# Pupillary manifolds: uncovering the latent geometrical structures behind phasic changes in pupil size

**DOI:** 10.1101/2024.05.23.595554

**Authors:** Elvio Blini, Roberto Arrighi, Giovanni Anobile

## Abstract

The size of the pupils reflects directly the balance of different branches of the autonomic nervous system. This measure is inexpensive, non-invasive, and has provided invaluable insights on a wide range of mental processes, from attention to emotion and executive functions. Two outstanding limitations of current pupillometry research are the lack of consensus in the analytical approaches, which vary wildly across research groups and disciplines, and the fact that, unlike other neuroimaging techniques, pupillometry lacks the dimensionality to shed light on the different sources of the observed effects. In other words, pupillometry provides an integrated readout of several distinct networks, but it is unclear whether each has a specific fingerprint, stemming from its function or physiological substrate. Here we show that phasic changes in pupil size are inherently low-dimensional, with modes that are highly consistent across behavioral tasks of very different nature, suggesting that these changes occur along pupillary manifolds that are highly constrained by the underlying physiological structures rather than functions. These results provide not only a unified approach to analyze pupillary data, but also the opportunity for physiology and psychology to refer to the same processes by tracing the sources of the reported changes in pupil size in the underlying biology.

**Graphical abstract:** 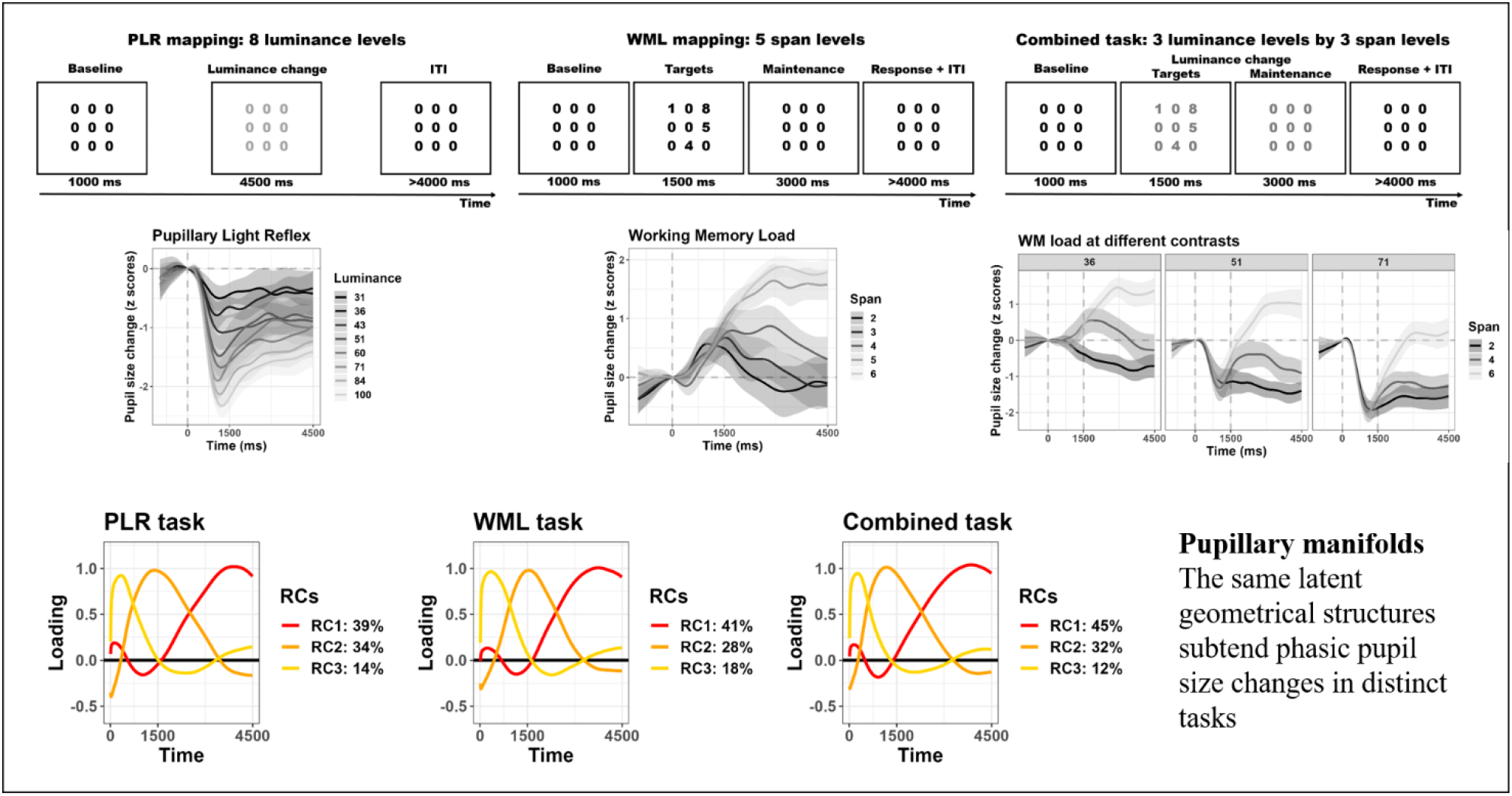

**Significance statement:** Phasic changes in pupil size are thought to reflect dynamic shifts between attentional states as instantiated by the locus-coeruleus noradrenaline system, and are crucial for adaptive behaviors. We found that the latent space of these changes is low-dimensional and remarkably similar across very different tasks, involving distinct cognitive processes. We therefore introduce the notion of pupillary manifolds as latent spaces that subtend the generative processes behind these changes. We suggest that manifolds arise due to hard constraints in the underlying physiological substrate – the relative balance between sympathetic and parasympathetic activity. In the framework outlined here, these mechanisms can be accessed and described directly, with only a handful of parameters, thus better informing computational modelling.

## Introduction

The pupils evolved primarily as a key tool for vision. Their core task is to manage the amount of light reaching the retina at any given moment, depending on the environment, as to optimize visual acuity ^1–3^. The nature of pupillary responses to light, i.e. pupil constriction, is therefore largely reflexive. However, pupil dynamics that do not have a strict environmental explanation (in terms of light levels) also exist; decades of research have ascribed them to a plethora of distinct cognitive processes spanning attention, emotion, working memory load, and executive functions more generally ^3–11^. Even the most fundamental **Pupillary Light Reflex (PLR)** is not completely impervious to cognitive, top-down modulations ^12^. For example, the PLR is increased, and the pupils constrict more, whenever stimuli are attended, even if covertly ^13–16^. The pupils can dilate, instead, for all sorts of arousing, demanding tasks and stimuli ^17–21^, often included under the same umbrella term of “psychosensory” modulators ^3^. The dilation response is notoriously sluggish, and is only seen several hundreds of milliseconds after the trigger event and the corresponding early orienting reflex, which also yields effects that are protracted in time. This means that pupillary responses that are functionally distinct typically overlap, on the one hand, and that searching for timepoints with significant differences in pupil size between experimental conditions is questionable at best, on the other hand. Regardless of the chosen analytical procedure, it is clear that as soon as one timepoint shows a significant difference this is the result of fully-blown latent processes and their interaction. Here we therefore turned to dimensionality reduction techniques to unveil these latent, unobserved processes and characterize them. The question was whether functionally distinct pupillary signatures can be mapped onto a low-dimensional space, and if so whether this space reflects the specific function mapped by the task (and experimental context) or rather unspecific physiological constraints.

## Results

We started with mapping the pupillary responses of 20 healthy human participants to stimuli changing (slightly) in luminance. We asked participants to simply view a matrix of numbers superimposed on a black background: nine “zeros” were arranged in three rows and intermittently changed their shades of gray so that we could map phasic responses to 8 different luminance levels (**Figure 1A**). We found the well-known PLR, consisting in a rather quick constriction of the pupils which reaches its peak within 1/1.5 seconds and then gradually recoups to a new baseline diameter (**Figure 2A**). We performed dimensionality reduction first with Principal Components Analysis (PCA) to obtain features that maximally summarize the variability in the data. We found that one component (PC1), i.e. one score value per trial, was sufficient to represent 79% of all the data (3 scores represented 95%, **Supplementary Figure 1**). The data are also well summarized by the eigenvector of PC1 (**Figure 2B**), describing the relevance of each timepoint to the component, which clearly depicts the “shape” of the PLR. While this is not surprising, because pupil traces are strongly autocorrelated, it remains remarkable how compact such a rich dataset and physiological measure can become. Having to cope with only a handful of values per trial is more manageable for most uses. Furthermore, the odds that components start to represent noise, instead of genuine signal, increase with their number; in this sense resolving to use few, quintessential features allows one to probe the primitive shape of the process at hand, e.g. the PLR, directly, without a priori assumptions. In **figure 2C** we show that this approach is also powerful in that PC1 clearly separates all the 8 different luminance levels administered in the task (F_(1, 527.75)_= 144.77, p <.001), thereby corroborating the notion that it could represent a latent dimension along which the PLR happens with different strength.

**Figure 1:**
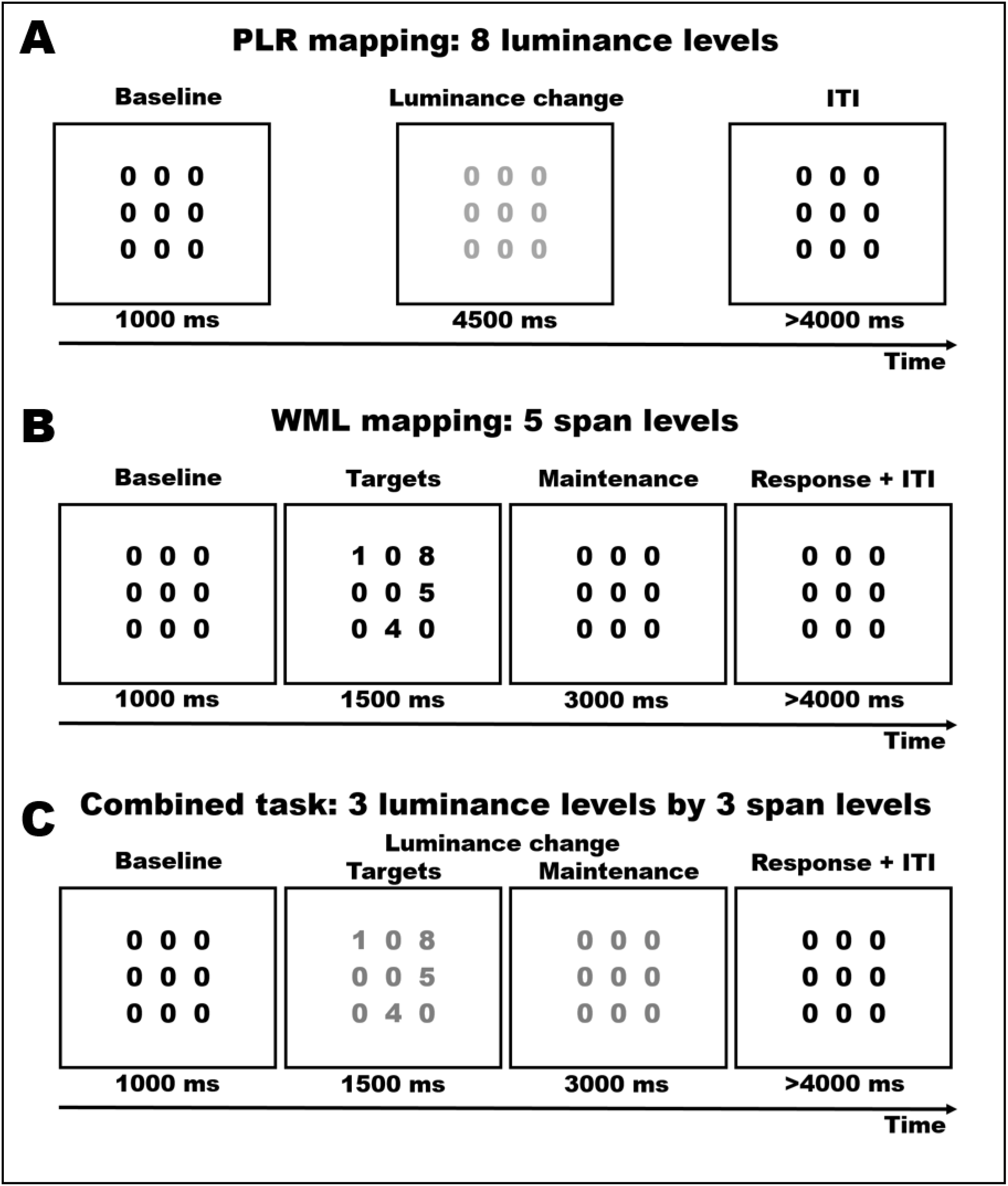
experimental procedures. We started with mapping the pupillary reflex to light (PLR, **panel A**). We asked participants to passively view a matrix of gray numbers superimposed on a black background. In each trial, the luminance of the numbers changed randomly to one of 8 brighter grayscale values, whereas the numbers themselves never changed. In **panel B**, on the other hand, we mapped pupil size changes to different working memory load (WML) conditions. While the luminance of the stimuli never changed, 2 to 6 numbers were selected randomly to replace the zeros, in random positions in the matrix. These target numbers remained in position for 1500 ms before being replaced by the baseline (nine “zeros”) for additional 3000 ms; numbers thus had to be kept in memory throughout this duration, until a response was solicited by an auditory sound. Finally, **panel C** depicts the combined task, in which both luminance levels and cognitive load were manipulated in a 3 x 3 design. Here, either 2, 4, or 6 randomly selected numbers appeared in random positions in the matrix, for 1500 ms, and then disappeared; concurrently, the luminance of the numbers changed randomly to one of 3 brighter values.

**Figure 2:**
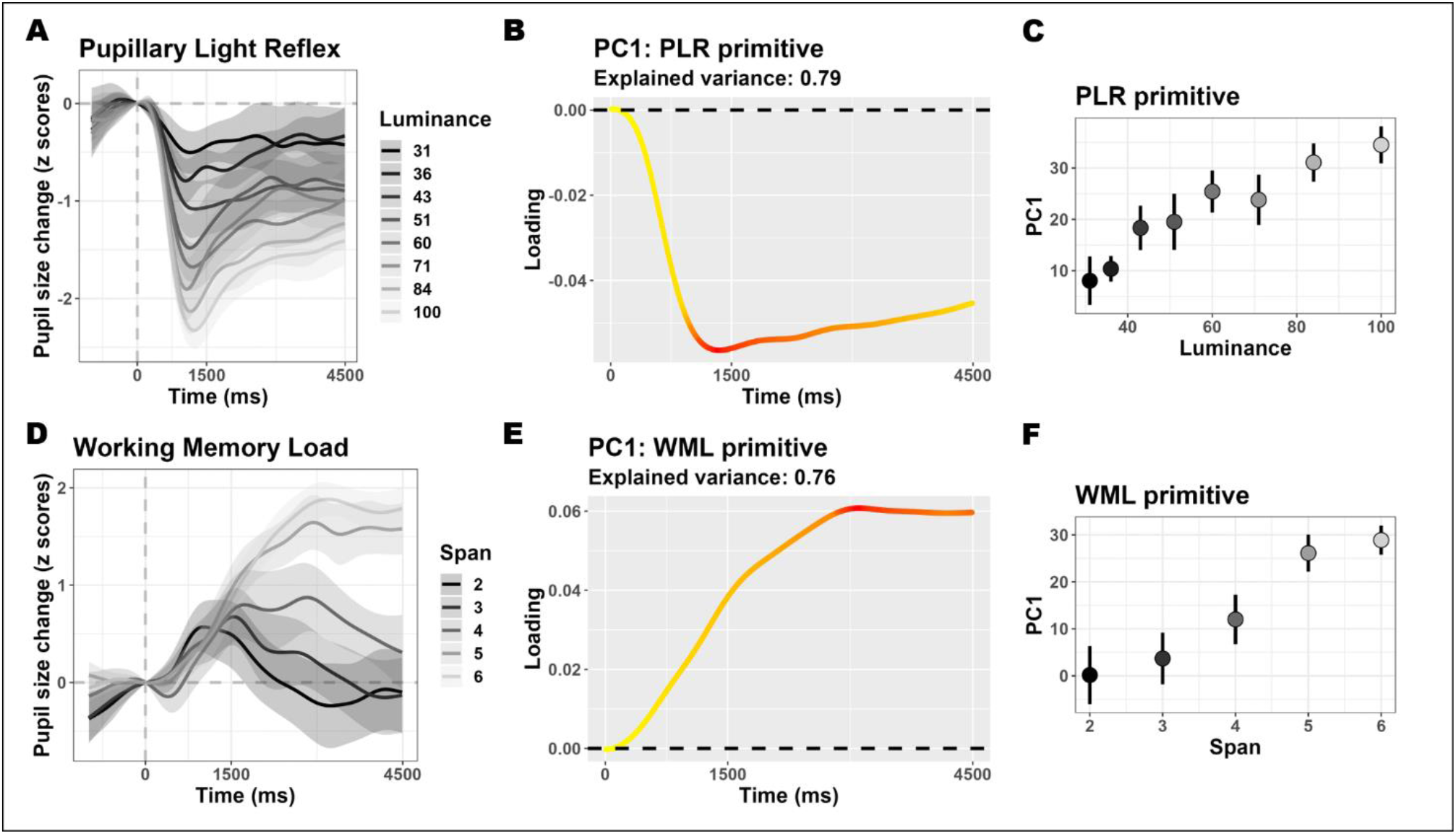
mapping pupillary responses to light and cognitive load. **Panels A and D** depict the time course of pupil size changes in the respective tasks; the shaded areas depict the 95% confidence interval. The first principal component from each task could explain a sizeable portion of the overall variability in the data. The respective eigenvectors are depicted in **panels B and E**, and capture the primitive shapes of the processes at hand. In these panels, the color scale ranges linearly from yellow\smallest weight to red\largest weight. Both principal components’ scores map efficiently a latent space along which the respective processes happen with a distinct strength. This mapping is depicted in **panels C and F** (error bars represent 95% confidence intervals).

We then moved to mapping the pupillary responses to changing **Working Memory Load (WML)**. In this case, the presented numbers did not change their luminance. Instead, for 1.5 s, the matrix was populated with 2 to 6 numbers chosen randomly and appearing in random positions (**Figure 1B**). Participants had to scan the matrix first, and then retain the numbers in working memory for an additional 3 s before providing their response, following the presentation of an auditory cue. We thus adapted a rather classic working memory span task in order to avoid the sequential presentation of stimuli, in consequence of which storing in memory (and pupil dilation) would presumably occur at different moments in different load conditions ^22^. The adaptation was effective because: behavioral performance declined with increasing WML (**Supplementary Figure 2**); pupillary responses clearly showed robust dilation as a function of WML (**Figure 2D**, F_(1, 18.66)_= 137.82, p <.001). As observed for the PLR mapping, this task also provided a dataset composed of very few latent dimensions. One principal component could account for 76% of the variability, 3 components about 93% (**Supplementary Figure 1**). The first component clearly distinguished different WML levels (**Figure 2F**), and its eigenvector (**Figure 2E**) suggests the primitive shape of WML is a slower, sustained dilation during the course of the trial and memory maintenance.

Next, we administered a task in which both dimensions changed in a 3×3 design, that is, both luminance and the amount of numbers to memorize were manipulated (**Figure 1C**). Both main effects were evident in pupillary recordings and resembled those described in the previous tasks (**Figure 3A**), also behaviorally (**Supplementary Figure 3**). Once again, the dataset was inherently low-dimensional, and the first component could account for 73% of the overall variability (3 components for 94%, **Supplementary Figure 1**). In this case, PC1 tracked well both features (Luminance: F_(1, 64.91)_= 28.62, p <.001; Memory load: F_(1, 21.97)_= 41.18, p <.001; **Figure 3C**) but did not discriminate between the two. Indeed, the eigenvector of PC1 (**Figure 3B**) fell almost perfectly midway between the PLR and WML fingerprints described above (**Figure 3D**). While, again, this remains a handy, effective way to summarize a dataset without too many arbitrary choices (e.g., a given time window) or assumptions (e.g., the shape of the pupillary function), it remains to be assessed whether this low-dimensional space maps onto distinct functional processes.

**Figure 3:**
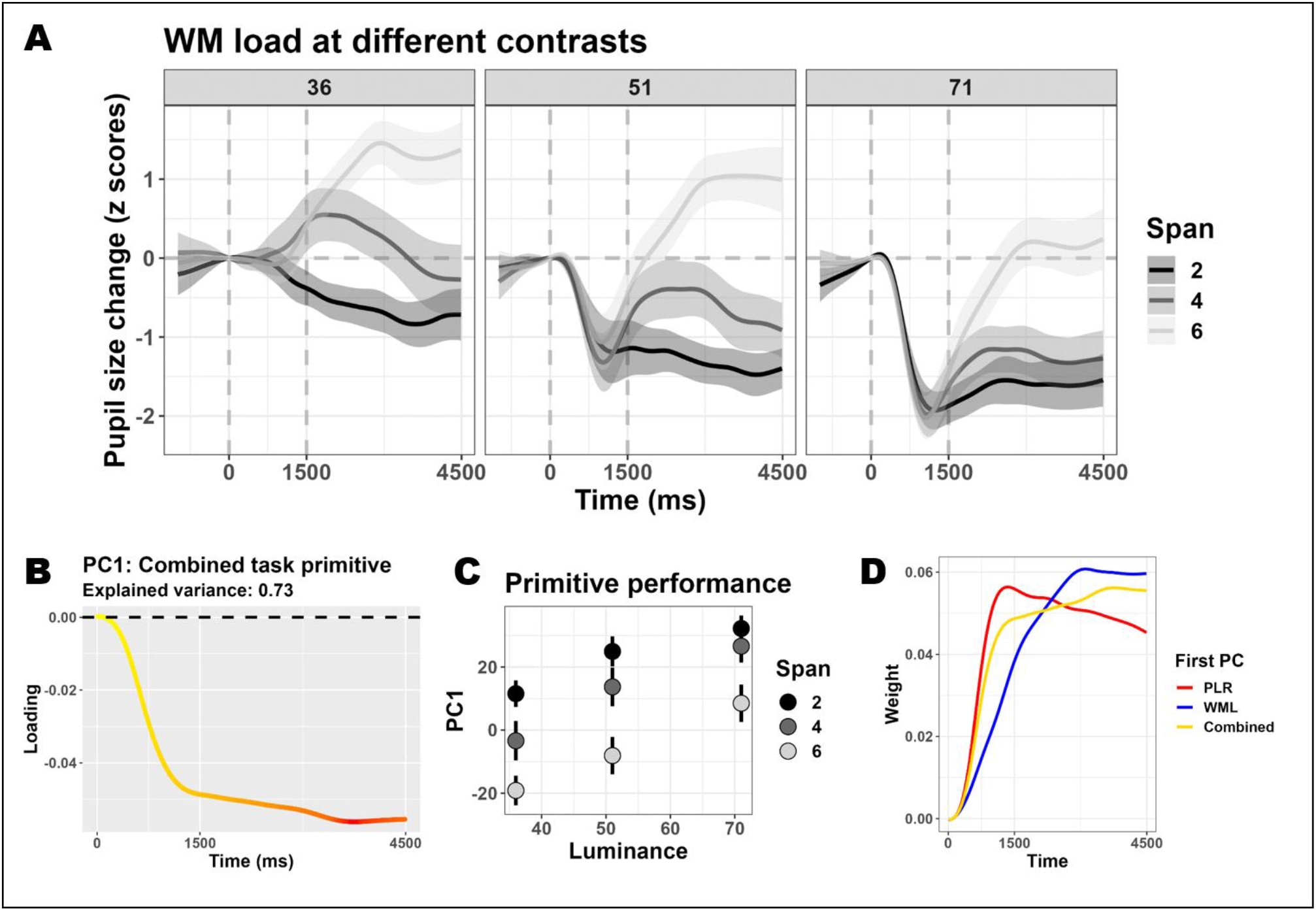
mapping pupillary responses to changes of both light and cognitive load. **Panel A** depicts the time course of pupil size changes as a function of different working memory load conditions (colors) and luminance levels (facets); the shaded areas depict the 95% confidence interval. This combined task was also characterized by a very low dimensionality. The eigenvector of the first component is depicted in **panel B** (the color scale ranges linearly from yellow\smallest weight to red\largest weight). The scores obtained from this component mapped very efficiently both dimensions (**panel C** depicts the mean scores, error bars represent 95% confidence interval). **Panel D** depicts the eigenvector (as in B) together with those obtained when mapping PLR and WML in isolation (as in **2B** and **2E**, though the sign was flipped for the former).

We therefore turned to non-orthogonal approaches with the aim of identifying latent processes that are potentially correlated as well as more interpretable. We focus in particular on promax-rotated PCA (rPCA), in keeping with previous suggestions ^23–25^. We choose to isolate k= 3 components following these studies and because components above three accounted for less than 3% of overall variability each in PCA. These analyses yielded three rotated components which were remarkably similar across the three different tasks, albeit accounting for a slightly different share of the overall variability (**Figure 4A**). Despite that, and despite oblique rotations are often non-optimal to account for the largest variability sources in the data, three components were sufficient to account for most of the phasic pupillary dynamics in all tasks (>87%). Because the three tasks vary substantially in terms of their requirements and pupillary signatures, these components are unlikely to reflect task-specific features. Rather, since this structure appears consistently across studies ^23–25^ and tasks of different nature, they are more likely to reflect underlying physiological constraints of the signal (changes in pupil size) rather than function (either pupil constriction to light or dilation to memory load). Indeed, both luminance and memory load in the combined task were equally well-tracked by the first two rotated components, which could discriminate both (Luminance: F_(1, 55.78)_= 27.21 and p <.001 for RC1, F_(1, 3.58)_= 23.96 and p =.011 for RC2; Memory load: F_(1, 26.52)_= 43.49 and p <.001 for RC1, F_(1, 302.22)_= 50.92 and p <.001 for RC2; **Figure 4B**). The exception was RC3: this component showed up consistently across tasks, with the earliest latency, and explained the smallest share of the overall variance. RC3 was sensitive to changes in luminance in the combined task (F_(1, 72.74)_= 13.88, p <.001; **Figure 4C**) and in the PLR mapping (**Supplementary Figure 4**), but not cognitive load in either the combined or WML mapping tasks (F_(1, 22.09)_= 0.63, p =.437; **Figure 4C** and **Supplementary Figure 5**).

**Figure 4:**
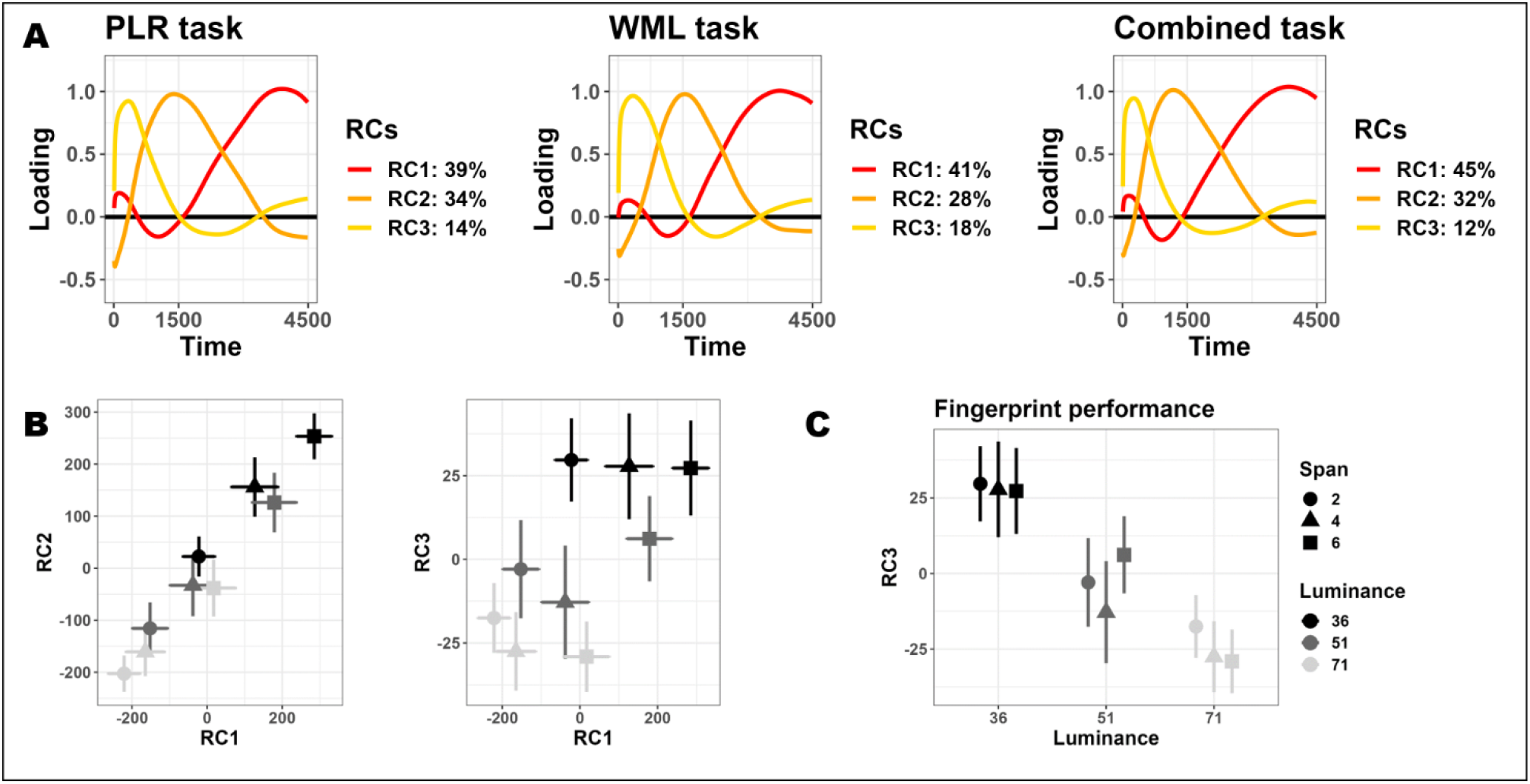
latent pupillary dynamics unveiled by oblique factor analysis. Regardless of the task and cognitive processes at hand, we found three remarkably similar structures behind the data (**panel A**). These three components could explain a large portion of the data and mapped efficiently both PLR and WML. **Panel B** depicts biplots with scores from pairs of the three components obtained in the combined task; plotted are mean scores (error bars are 95% confidence intervals). The first two components mapped constriction to light and dilation to cognitive load equally well. On the other hand, one component, RC3, mapped reflexes to light specifically (**panel C**).

## Discussion

In this study we decided to map pupillary reflexes to light, which are far more robust than, for example, the modulations of the PLR that may be caused by attention ^13,14^. Furthermore, these reflexes were contrasted against psychosensory modulations of pupil size induced by working memory load. This means that we were in a rather favorable position to maximize the chances to isolate processes that are both functionally and physiologically distinct, considered that their neural generators are very different ^12,26,27^. The point was not as much about showing that changes in pupil size are inherently low-dimensional – this is expected in light of the strong autocorrelation of the signal – but rather how well the latent, low-dimensional space separates distinct functional processes.

We did find one component (RC3) that appeared to map luminance changes but not pupil dilation due to cognitive load. This component accounted for a small share of the overall variance in the data and loaded especially on the earliest timepoints, which may suggest it refers to the PLR specifically. However, it is striking that this component was still recovered as such in a task only probing memory load, and even though it did not discriminate between key conditions; this points to an underlying latent structure that remains deployed, possibly silently, regardless of the task at hand (e.g., a primarily parasympathetic component). Overall, indeed, we found that the components recovered from very different tasks and pupil traces were very similar; this by itself speaks about the presence of common constraints in these traces. Moreover, the same components, beyond RC3, could track both reflexes to light and mental effort, which lead to signature changes in pupil size. The fact that these functionally different processes can be quantified very well by few components with a similar shape points again to the existence of constraints of a different nature. For example, a few authors have suggested, based on pharmacological interventions or the manipulation of environmental light levels, that pupillary dilation to mental effort may be composed of two waves ^28,29^: the first, predominant in bright environments, would reflect the cortical inhibition of the parasympathetic efferent pathway; the second, occurring later in time and explaining most of the dilation, would represent instead a primary sympathetic component. One should be weary in over-interpreting oscillatory patterns in PCA ^30^. However, this account fits well with our findings, and could very well outline the main constraints imposed on pupil size, beyond its autocorrelated nature. Contextual variables may tap onto the dynamic continuum between the different branches of the autonomic nervous system, thereby causing pupil size to change along low-dimensional manifolds that reflect this balance accurately, but much more efficiently. Clearly, the blanket is too short to separate neatly distinct functional processes, as increased sympathetic activity implies, with the passing of time, decreased parasympathetic tone. Still, here we show that changes in pupil size are not different from much richer physiological signals in their potential to be mapped onto low-dimensional, interpretable spaces ^31,32^. The notion of neural manifolds has enriched greatly the current debate around the neural control of movement or other cognitive processes; we surmise that the notion of pupillary manifolds could, likewise, provide fruitful ground for cognitive and computational pupillometry. The possibility to infer latent, generative processes behind the observed signal is of particular interest. These processes may sometimes be hard to see from the data, and yet constitute the source biological signal that explains most of the subsequent changes in pupil size; for most uses, this is therefore the signal of interest. Ultimately, the concept of pupillary manifold provides a tool to delve deeper into the physiological origin of phasic changes in pupil size. These changes have been clearly implicated in finely-tuning behavior toward maximal utility ^33–35^, so that accessing and better characterizing their fingerprint on pupil size represents a very appealing challenge, one which would contribute harmonizing the terminology between psychology and physiology. Last but not least, this feature points to the opportunity of a unified approach to analyze pupillary data via dimensionality reduction, for which we provide a toolbox (https://github.com/EBlini/Pupilla). This approach not only shields from more or less arbitrary assumptions about the latency (e.g., time windows) or the shape (e.g., pupillary response function) of pupillary dynamics (which open an entire multiverse of analytical choices, ^36^): it could feed much more efficiently physiologically-informed computational models of human learning and decision-making.

## Methods

All materials, raw data, and analysis scripts for this study are available through the Open Science Framework website: https://osf.io/dkpcs.

The core functions for preprocessing and analysis are available through GitHub: https://github.com/EBlini/Pupilla.

### Participants

Twenty participants, all students from the University of Florence, took part in this study (17 females, M= 25.1 years, SD= 6.95 years). Inclusion criteria were normal (or corrected-to-normal) vision, and no history of neurological, psychiatric, or sensory disorders. The experimental procedures were approved by the local ethics committee (Commissione per l’Etica della Ricerca, University of Florence, July 7, 2020, n. 111). The research was carried in accordance with the Declaration of Helsinki, and informed consent was obtained from all participants prior to the experiment.

### Procedures

The participants were tested in a dimly lit, quiet room, their head comfortably resting on a chinrest. They faced a remote infrared-based eye-tracker (EyeLink 1000, SR Research Ltd.), at a distance of approximately 57 cm from the screen. Each session started with a 15-points calibration of the eye-tracker, which was then set to monitor participants’ pupil size continuously at a 500 Hz sampling rate. The open-source software OpenSesame ^37^ was used to present the stimuli, along the procedure outlined below and depicted in **Figure 1**.

### PLR mapping

We asked participants to passively view a matrix of gray numbers superimposed on a black background (0 on the grayscale, 0.8 cd/m^2^, **Figure 1A**). The numbers were nine “zeros” encompassing about 3°, written in gray (30 on the grayscale). In each trial, numbers were presented as such for 1000 ms during the baseline phase. Then, for 4500 ms, the luminance of the numbers changed randomly to one of 8 grayscale values (31, 36, 43, 51, 60, 71, 84, 100). The 8 levels were chosen to be approximately linearly arranged on the logarithmic space. In absolute terms, these luminance values mapped roughly linearly in the range between 3.5 and 27 cd/m^2^ (note that this reflects the color of the font, not the local luminosity of stimuli). After this phase of interest, the numbers returned to the original intensity value and were presented for 4000 to 5000 ms (with a uniform jitter), in order to allow the pupils to return to their baseline size. There were 48 trials, 6 for each luminance level.

### WML mapping

The stimuli during the baseline and ITI phases were identical to the PLR task, though their color intensity (on the grayscale) was slightly higher (51); this was because the focus was pupillary dilation to working memory load, and this minor adaptation allowed for more room to observe changes in this direction. In this task, the numbers presented in the phase of interest did not change their luminance. Instead, for 1500 ms, the matrix was populated with 2 to 6 numbers chosen randomly and appearing in random positions (**Figure 1B**). Participants had to scan the matrix first, and then retain the numbers in working memory for an additional 3000 ms before providing their response. During this time, the numbers returned identical to the baseline (nine “zeros”). Once this phase ended, an auditory cue prompted participants’ responses, which was then scored manually by the researcher. This task was composed of 40 trials, 8 for each of the 5 levels of cognitive load.

### Combined task

In this task, both dimensions (luminance and cognitive load) changed in a 3×3 design (**Figure 1C**). The stimuli during the baseline and ITI phases were identical to the PLR task. Then, either 2, 4, or 6 randomly selected numbers appeared in random positions in the matrix; concurrently, the luminance of the numbers changed randomly to one of 3 brighter values (36, 51, 71 on the grayscale). After 1500 ms, the matrix returned to being composed only by zeros, though luminance manipulation remained in place for an additional 3000 ms. The participants’ response was solicited by an auditory cue (as in the WML task above), and manually recorded by the experimenter. This task was composed of 90 trials, 10 for each of the 9 conditions given by the 3 x 3 design (Span by Luminance); it was administered in two blocks to avoid participants’ excessive fatigue and distress.

### Data processing

We only retained pupil size measured during fixations, as identified by the eyetracker (i.e., measures of pupil size during blinks and saccades were discarded). We did not further correct pupil size by regressing out the position of the gaze on the screen. However, we don’t have a reason to suspect this may constitute a bias because the eyes were highly constrained to central vision (< 3°), and whenever numbers beyond zero were presented (i.e., WML and combined task) they appeared in random positions of the matrix. In addition, we further used a velocity-based criterion to identify likely artifacts; each gap in the traces was then extended by 20 ms both before and after the artifacts. Trials in which more than 40% of the data were missing were excluded, and the gaps in the remaining ones were linearly interpolated. Traces were then smoothed through cubic splines. Next, traces were down-sampled to 10 ms epochs by taking the median pupil diameter for each time bin. In order to better cope with inter-individual differences we then z-transformed pupil diameter values separately for each participant and task ^15,19,38^. With normalization, a value of 0 represents the subject-specific mean pupil diameter and, regardless of baseline values, scores represent the relative pupil size expressed as a fraction of the overall participant’s variability. This is useful to account for the fact that the same amount of relative dilation or constriction (e.g., 0.1 mm) has very different meanings in participants with small vs. large baseline pupil size. Trials starting with extreme baseline values (exceeding 2 sd from the mean of the baselines of all trials) were discarded. Finally, all series were realigned by subtraction to a baseline value set to be the median of pupil size in the time window located ±250 ms from the onset of the phase of interest. Overall, data cleaning led to discard an average of: 9.5% (SD= 6.5%) trials per participant in the PLR mapping task; 11.2% (SD= 12%) trials per participant in the WML mapping task; 13.1% (SD= 14%) trials per participant in the combined task. Overall, the final pupillometric results were in keeping with classical findings regarding the pupillary light reflex ^3^ and psychosensory modulations (i.e., by cognitive load) ^22^.

### Dimensionality reduction

We used R version 4.2.3 ^39^. As a first step we used temporal PCA through the ‘prcomp()’ function, which uses single value decomposition. We did not further centered and scaled the data matrix, which included data from all participants and trials, because pupil size data were normalized and baseline-corrected beforehand, as outlined above. Temporal PCA has been used before to describe the major component(s) of pupil traces, including that of the PLR ^14^. Previous studies have also attempted to link different components to different functions, for example a sustained response versus more residual responses (e.g., transient responses, cognitive variables, accommodation-related phenomena) ^14,40^. However, in the case of tasks that elicit multiple, heterogeneous processes (e.g., both the PLR and psychosensory-dilation effects) this becomes less viable: PCA attempts to describe the original data in the most compact way possible, but this efficiency in data reduction sometimes hampers the interpretability of the eigenvectors as latent constructs. In the context of this study, this approach resulted in the eigenvector of the combined task being a combination of the eigenvectors recovered in the PLR and WML tasks. This could be a convenient analytic tool to draw statistical inferences, but it appears clear that a functional interpretation of the component would be unwarranted in this case. Rather, rotations can be used to transform the best PCA solution into alternative ones which, at the cost of being less efficient in terms of data reduction (components are not “principal” anymore), yield eigenvectors that are more easily interpretable. Rotations, such as *promax* used here, can be oblique, which allows components to correlate instead of being orthogonal – often a more realistic and biologically sound scenario. Note that here we followed the ‘*psych*’ package ^41^ and rescaled the eigenvectors by the square root of the eigenvalues, which makes them more comparable to factor loadings more typical of factor analysis.

### Analyses

The following packages greatly eased our work: psych (v 2.4.1, Revelle, 2024); dplyr (v 1.1.4, Wickham et al., 2023); ggplot2 (v 3.4.4, Wickham, 2016); lme4 (v 1.1-35.1, Bates et al., 2015); afex (v 1.3, Singmann et al., 2023).

When analyzing pupil traces, we followed ^46^ by using crossvalidated Linear Mixed Effects Models (LMEM) through the lme4 package for R ^44^. In this approach all trials from each participant are assigned deterministically to one of 3 folds. Two folds are circularly used as the training set; here, intercept-only LMEMs are performed for each timepoint, and the timepoint having the peak t-value (for each fixed effect or interaction) is then used in the test set to confirm the overall consistency of the target factor across folds. This approach is computationally efficient and very powerful in suggesting the presence of a consistent experimental effect *somewhere* along the time course of the trials. In order to enhance the precision in identifying a temporal cluster for any given effect, we additionally scored a consensus between folds, as timepoints in which all folds presented t-values above |2|; timepoints in which this rather stringent, albeit arbitrary criterion was met were thus identified as a temporal cluster ^38^.

We then analyzed the scores obtained from dimensionality reduction techniques. The scores were obtained, for each trial, through the ‘predict’ method of the relevant function. For inference, we also turned to type 3 LMEMs with p-values given by the Satterthwaite approximation as implemented in afex (v 1.3, Singmann et al., 2023). In this case we could always set the matrix of random effects to be the maximal one ^47,48^, that is one including random slopes for all fixed effects and their interactions.

### PLR mapping

There was a significant main effect of Luminance (*t*_(848.78)_= -14.73, *p*<.001; peaks around 1.1 s). A clear consensus for this effect was found starting from 400 ms and lasting until the end of the trial (**Figure 2A**).

Luminance could also predict the scores of the first component (**Figure 2C**): F_(1, 527.75)_= 144.77, p <.001.

### WML mapping

There was a significant main effect of working memory load (*t*_(690.75)_= 14.91, *p*<.001; peaks between 3 and 3.5 s. A clear consensus for this effect was found starting from 1.5 s and lasting until the end of the trial (**Figure 2D**).

Working memory load could also predict the scores of the first component (**Figure 2F**): F_(1, 18.66)_= 137.82, p <.001.

### Combined task

In the combined task, both main effects were significant (**Figure 3A**). There was a significant main effect of Luminance (*t*_(1541.64)_= -9.19, *p*<.001; peaks around 1 s and consensus from 440 ms up to 4.2 s). There was also a main effect of memory load (*t*_(1543.15)_= 7.69, *p*<.001; peaks between 2.8 s and 3 s, and consensus from 1.2 s onward). However, there was no interaction between the two (*t*_(1542.05)_= - 1.09, *p=*.275).

Both main effects were also significant when analyzing the first principal component scores (Luminance: F_(1, 64.91)_= 28.62, p <.001; Memory load: F_(1, 21.97)_= 41.18, p <.001; **Figure 3C**). In addition, however, there was a significant two-way interaction between the two (F_(1, 52.24)_= 4.34, p =.042). Linear contrasts showed a trend, in the condition with the brightest luminance, for a reduced effect of working memory load, especially between 2 and 4 digits (**Figure 3C**).

The effects above were almost identical when analyzing the first fingerprint (rotated component RC1) in that both main effects (Luminance: F_(1, 55.78)_= 27.21, p <.001; Memory load: F_(1, 26.52)_= 43.49, p <.001) and their interaction (F_(1, 58.51)_= 4.3, p =.042) were significant. The same holds for the second fingerprint, RC2 (Luminance: F_(1, 3.58)_= 23.96, p =.011; Memory load: F_(1, 302.22)_= 50.92, p <.001; Luminance by Load: F_(1, 31.48)_= 4.74, p =.037). Results are depicted in **Figure 4B**.

However, RC3 was found to be specific for Luminance (F_(1, 72.74)_= 13.88, p <.001), but not working memory load (F_(1, 22.09)_= 0.63, p =.437) or their interaction (F_(1, 34.79)_= 0.9, p =.348). RC3 is depicted in both **Figure 4B** and **Figure 4C**.

## Acknowledgements

This publication was produced with the co-funding of the European Union - Next Generation EU, in the context of The National Recovery and Resilience Plan, Investment 1.5 Ecosystems of Innovation, Project Tuscany Health Ecosystem (THE), ECS00000017. Spoke 3. CUP: B83C22003920001.

## Authors Contribution

**EB:** Conceptualization; Methodology; Software; Formal Analysis; Investigation; Data curation; Visualization; Writing – original draft, review and editing. **RA:** Funding acquisition; Writing – review and editing. **GA:** Writing – review and editing.

## Supplementary figures captions

**Figure S1:**
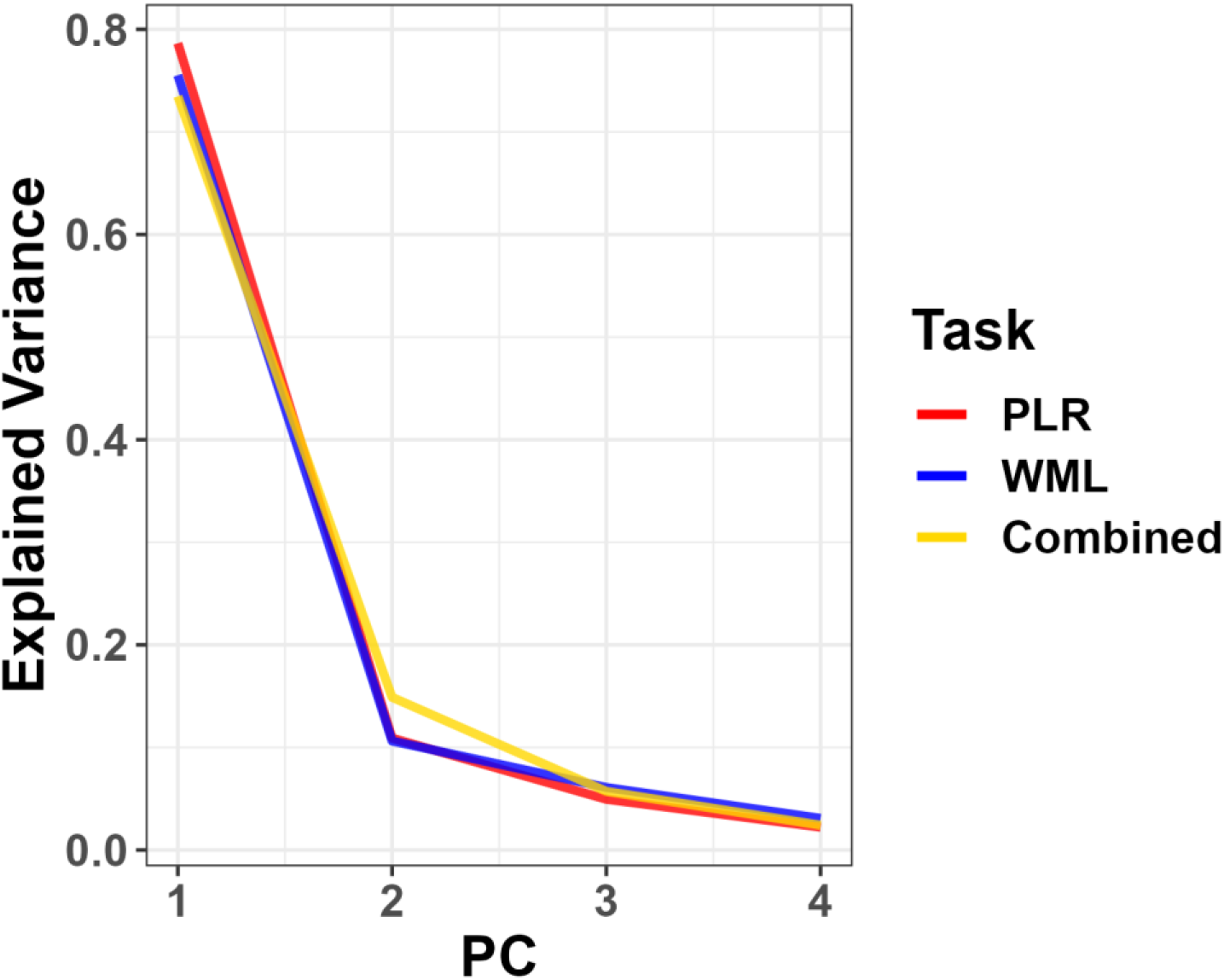
phasic pupil size changes are intrinsically low-dimensional. The figure depicts the scree plots from principal component analysis on the three tasks. Three components are generally sufficient to explain more than 88% of the overall variability in the generating data. Components above 3 account for less than 3% of the remaining variance.

**Figure S2:**
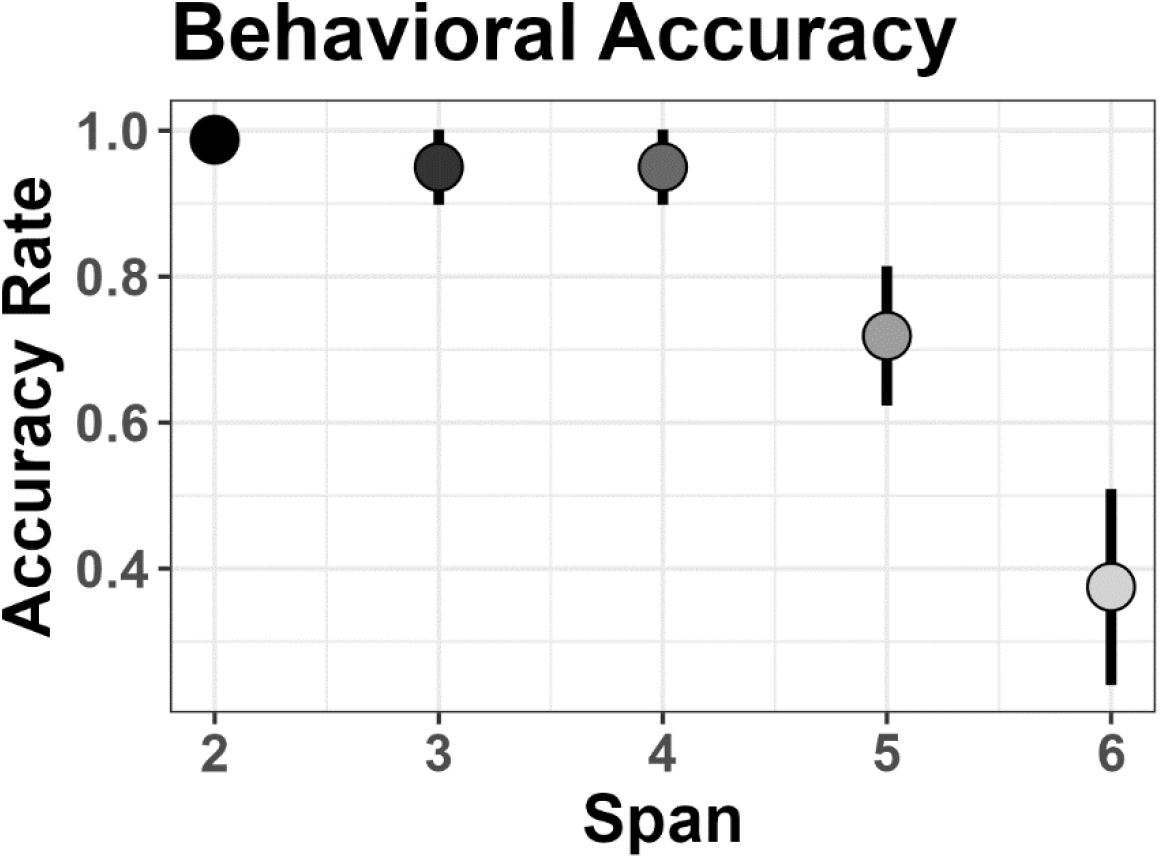
behavioral performance in the WML mapping task. Mean (95% confidence interval) accuracy in reporting the presented numbers as a function of cognitive load. Performance was near ceiling up until 4 digits, then decreased rapidly with 5 and, especially, 6 digits.

**Figure S3:**
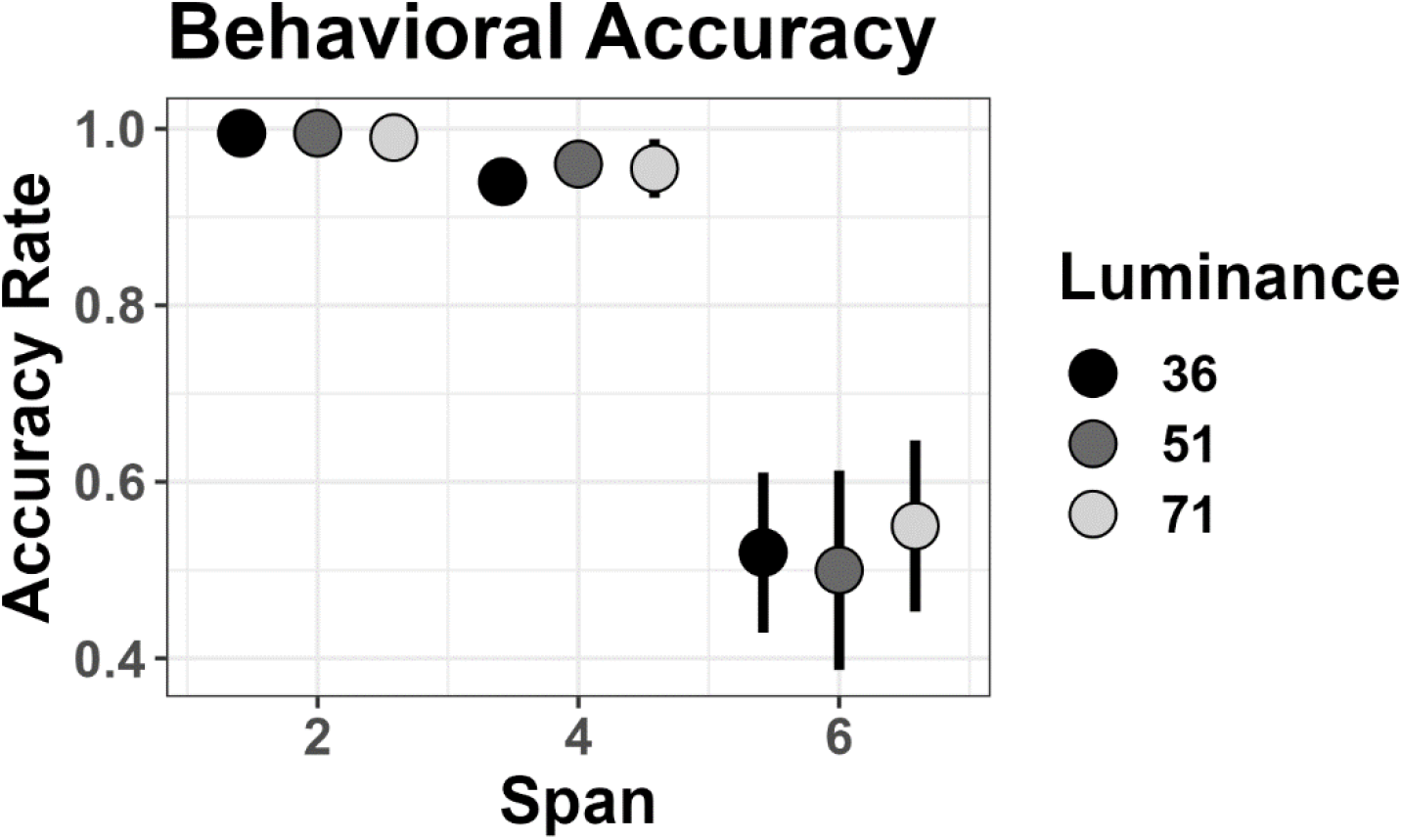
behavioral performance in the combined task. Mean (95% confidence interval) accuracy in reporting the presented numbers as a function of cognitive load and luminance level. Performance was near ceiling up until 4 digits, then sensibly decreased, regardless of luminance.

**Figure S4:**
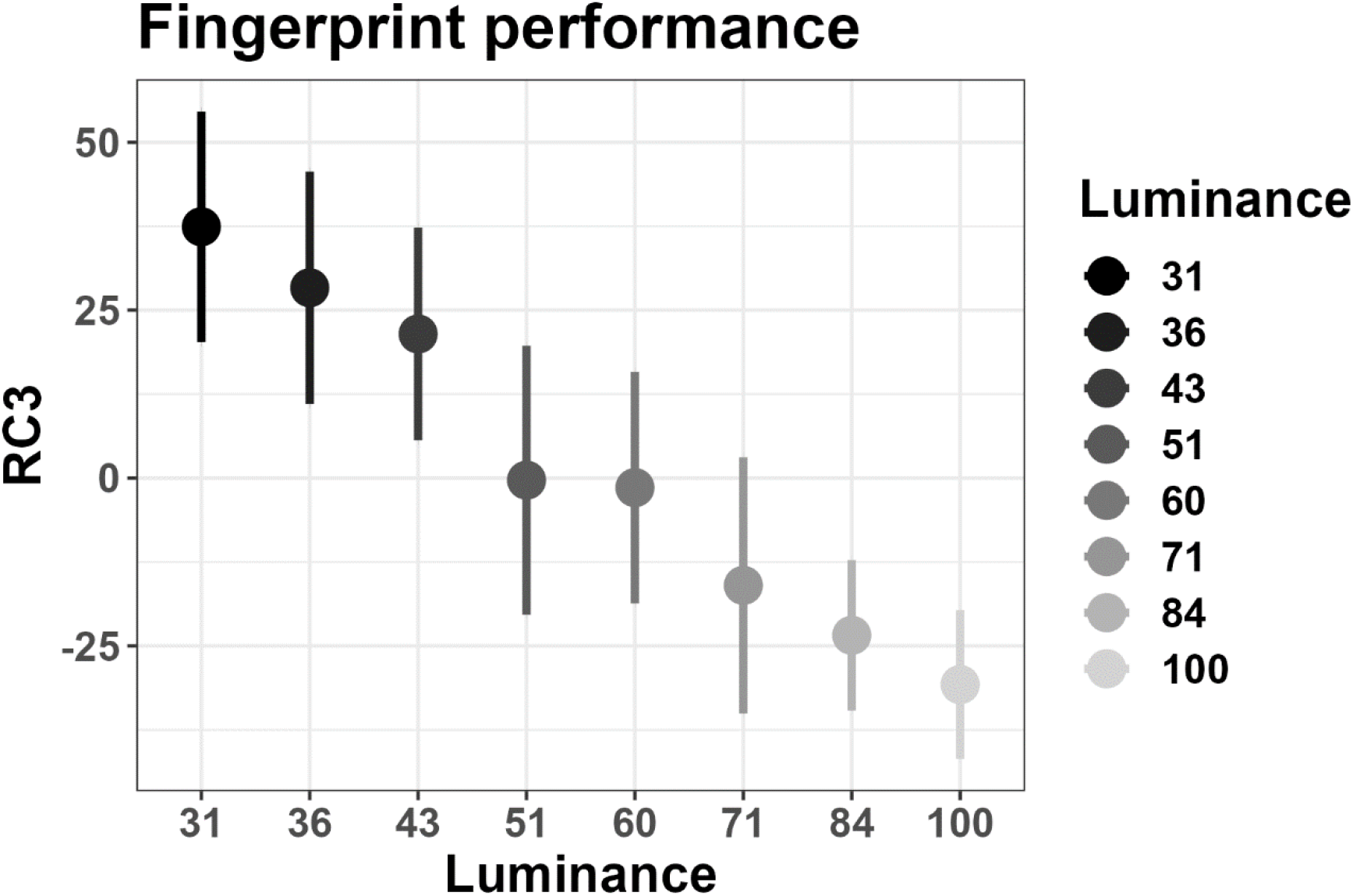
fingerprint component of reflexes to light. One rotated component, RC3, obtained from the PLR mapping task, mapped efficiently the entire luminance space on a latent dimension. Values depict mean scores (95% confidence interval).

**Figure S5:**
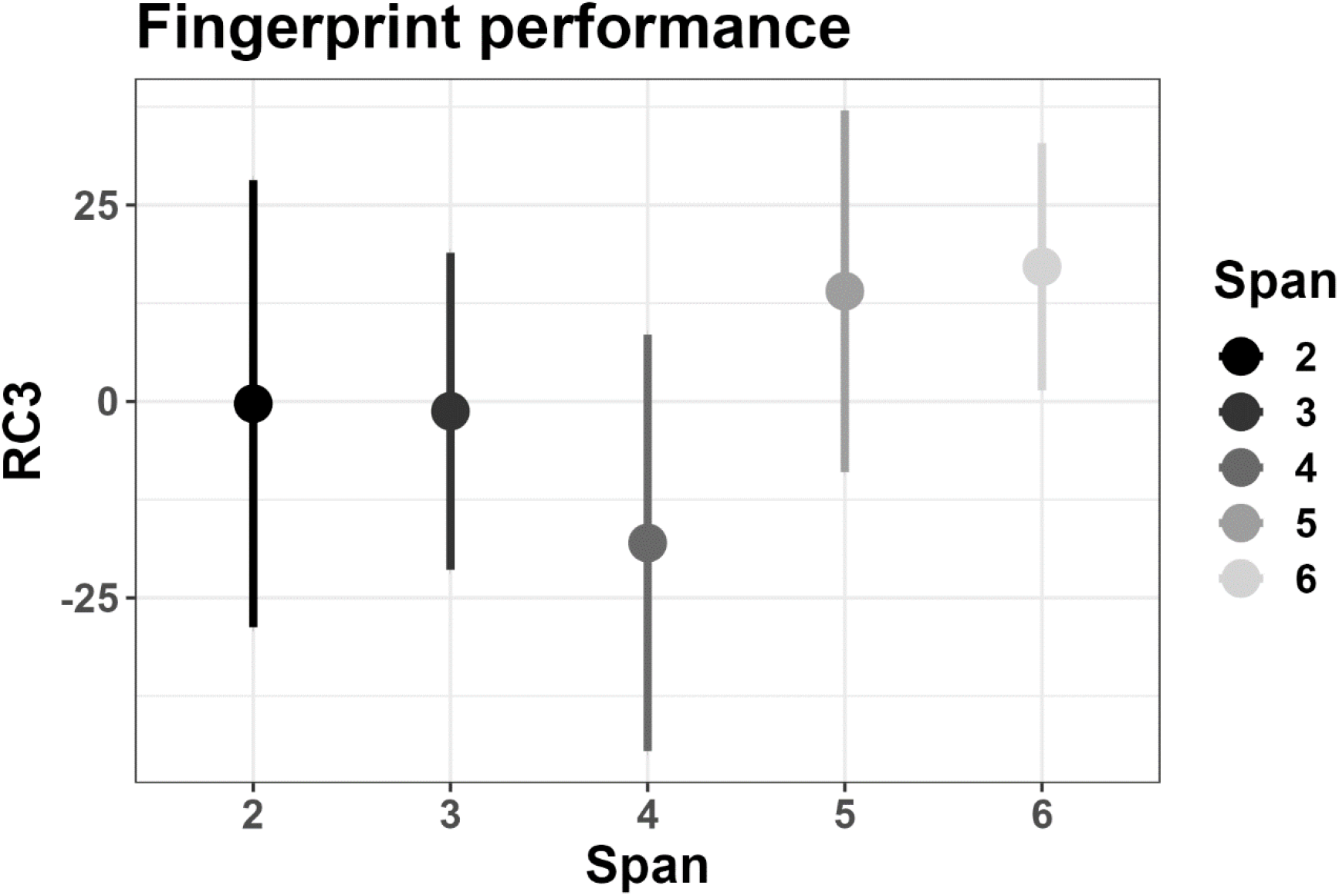
the fingerprint of reflexes to light do not map cognitive load. One rotated component, RC3, obtained from the WML mapping task, did not map distinct cognitive load conditions. Values depict mean scores (95% confidence interval).

